# The nightshift lowdown: can ants buffer climate change through shifts in vertical and temporal activity?

**DOI:** 10.64898/2026.02.17.702942

**Authors:** Lily Leahy, Brett R. Scheffers, Alan N. Andersen, Stephen E. Williams

## Abstract

Temperature fluctuations across space and time create a multidimensional thermal landscape within which organisms are exposed to local climates while conducting their daily activities. Tropical species are considered to be particularly sensitive to climate change, with narrow thermal safety margins - the buffer between operative and lethal temperatures. In tropical rainforests, however, species can hypothetically mediate thermal exposure via activity over two local thermal dimensions: vertical (ground-canopy) and temporal (day-night). Such spatiotemporal flexibility could protect species from elevated temperatures and improve thermal safety margins, but this mechanism has not been previously investigated. We test this hypothesis using rainforest ants at a warm lowland and cool upland site (100, 1200 m a.s.l.) in the Australian Wet Tropics. At lowland and upland sites, we quantified microclimate, foraging activity, community composition, and thermal ecology of ants across vertical and temporal dimensions. To assess spatiotemporal flexibility as a climate change mitigation strategy, we calculated thermal safety margins (TSM) as the difference between a species upper thermal limit (CT_max_) and mean activity temperature (T_e_). For each species in each of their spatiotemporal niches (ground-arboreal-day-night) we test whether shifting activity to cooler niches increases TSM using the hottest niches (arboreal and/or daytime) as a baseline. At both lowland and upland sites, ant species were highly stratified vertically, but the large majority (77 - 87.5%) were active both day and night, indicating widespread temporal generalisation. Shifting activity to cooler parts of the thermal landscape substantially improved TSMs: in the lowlands, species with arboreal diurnal foraging increased their TSM by an average of 4.4 °C (± 1.7 SE) by shifting to the ground and 6.7 °C (± 1.63 SE) by shifting to nocturnal foraging. Improvements were more modest in the uplands: arboreal diurnal foragers increased TSM by 2.1 °C (± 2.07 SE) and 2 °C (± 0.28 SE) for ground and nocturnal shifts respectively. We therefore demonstrate that foraging niche flexibility is an important climate-change mitigation trait and is most beneficial in the lowlands. Lowland diurnal canopy specialists, however, are most at risk. This represents a large proportion of tropical rainforest biodiversity, supporting previous hypotheses of lowland biotic attrition under climate change.

## Introduction

Tropical rainforest species are especially vulnerable to climate change because they typically have narrow thermal tolerances and limited elevational and geographic ranges (Huey et al. 2009). The rapid pace of climate change limits the capacity of many species to shift their ranges and there may be low potential for evolutionary adaptation (Stillman 2003, Moritz and Agudo 2013). The effects of rising temperatures, however, can be potentially mitigated *in situ* by behavioural responses that allow populations to exploit local thermal variability while remaining within their current ranges (Kearney et al. 2009, Sunday et al. 2014).

Spatial and temporal thermal gradients interact with each other to create climatic variability at the microscale (Potter et al. 2013). In forest systems, microclimatic variation is most pronounced during the day owing to the interaction between vegetation and solar radiation, whereas at night this microclimatic variation is less pronounced (Xing et al. 2023). Moreover, such local thermal gradients formed by habitat and forest structure are nested within broader scale climatic gradients of elevation and latitude. Species therefore exist in a multidimensional thermal landscape within which behavioural ecology determines thermal exposure while physiology determines sensitivity and persistence under local conditions (Bonebrake and Deutsch 2012, Huey et al. 2012).

In forests there are two thermal gradients along which species can be active, one relating to vertical space from ground to canopy (Leahy et al. 2021), and the other to the time of day (Kronfeld-Schor and Dayan 2003, Yusah et al. 2018). These local thermal gradients are spatial and temporal niche axes that represent a four-dimensional thermal landscape within which populations can manoeuvre to maintain optimal thermal exposure (Kronfeld-Schor and Dayan 2003, Basset et al. 2015). Species have evolved distinct activity patterns along these niche axes, ranging from specialist strategies, with limited spatiotemporal flexibility, to generalist strategies, with broad spatiotemporal flexibility (Cox et al. 2021, Leahy et al. 2021). For example, lemurs are generally cathemeral (active both day and night), but their level of diurnal activity depends on season, diet, group size, and level of anthropogenic disturbance (Donati et al. 2016). Flexibility can also be triggered by various environmental or biological cues; for example, amphibians in Panama shift downward from the canopy in the dry season and up again in the wet season when the canopy is more humid (Basham and Scheffers 2020).

Different temporal and vertical niche strategies may have evolved for a wide range of reasons, including maximising resource acquisition, avoiding predation and competition, as well as for thermal optimisation (Slavenko et al. 2022, Wong and Didham 2024). Whatever the case, in a warming world, generalist species could utilise this existing behavioural flexibility along spatiotemporal thermal gradients to buffer unfavourable temperatures *in situ* (Levy et al. 2019, Fredston et al. 2025). In contrast, specialised species that are restricted to the hotter parts of the environment will have less behavioural buffering capacity and therefore will be more vulnerable (Huey et al. 2012, McCain and King 2014).

Accurately characterising thermal exposure is essential to understanding species behaviour in localised thermal landscapes and how they might avoid temperatures close to their thermal limits (Potter et al. 2013, Storlie et al. 2014). Vertical microhabitats, time periods, and elevational sites may represent contrasting or similar thermal environments for small rainforest invertebrates depending on the degree of thermal overlap along these gradients. At different elevations, species may wish to buffer hot temperatures in the lowlands and cold temperatures in the uplands, leading to elevation-specific patterns of spatiotemporal movement (Bennie et al. 2014). In particular, we might expect more specialisation towards the warmer daytime temperatures and/or the canopy in cooler upland environments (Scheffers et al. 2013).

Rainforest ants are excellent organisms for investigating spatiotemporal patterns of foraging activity in relation to thermal exposure as they exhibit variation in activity along vertical, temporal, and elevational gradients and are highly sensitive to thermal conditions (Yanoviak and Kaspari 2000, Sanders et al. 2007, Houadria et al. 2015, Silva et al. 2019, Leahy et al. 2024). At local scales, ant thermal limits strongly relate to microclimate; canopy species tend to be thermal generalists and tolerate hotter temperatures than do ground or subterranean species (Baudier et al. 2015, Kaspari et al. 2015), while obligate nocturnal species can withstand colder temperatures than can diurnal species (Garcia-Robledo et al. 2018). Upland ant species can tolerate colder temperatures than lowland ants but tolerance of hot temperatures tends to be similar in lowland and upland ants (Bishop et al. 2017, Leahy et al. 2022).

Here, we use a multidimensional approach to investigate the foraging activity and thermal ecology of rainforest ants along vertical and temporal gradients and at contrasting elevations in the Australian Wet Tropics Bioregion. We ask whether flexibility in activity along spatiotemporal axes of verticality and time of day could confer resilience to climate change for ant communities and explore how thermal risks and opportunities change between lowland and montane environments. In both lowland and upland elevation sites, our aims are to: (1) quantify the thermal landscape by calculating thermal overlap of microclimatic temperature along vertical and temporal gradients, (2) investigate when and where species are active by characterising species composition and species activity over vertical and temporal gradients, (3) incorporate field active temperatures, ambient thermal exposure, and upper and lower thermal limits (hereafter, CT_min_ and CT_max_) to characterise the thermal ecology of ant communities, (4) estimate the increase in thermal safety margin (TSM) gained by foraging into cooler spaces and activity times. We use thermal safety margins as a metric of climate change risk defined as the difference between heat tolerance (CT_max_) and temperature exposure (T_e_). We use each of these lines of evidence to evaluate whether species could use flexibility in spatiotemporal activity to buffer rising temperatures in the rainforest and how these mitigation options change across elevation.

## Methods

### Study sites

Ants were sampled at two elevation sites at Mt. Lewis/Carbine mountain range (hereafter, Carbine), at a lowland, 100 m a.s.l. (Mossman Gorge; -16.470, 145.320), and upland, 1200 m a.s.l. (Mt. Lewis National Park; -16.510, 145.270) site. Sites were sampled from August – October 2019 and sampling was conducted only in the absence of rain (Leahy et al. 2022). The ant fauna at these sites has been previously described in (Leahy et al. 2022).

### Ant surveys

At each elevation site, we established five plots at least 50 m apart and selected the largest tree per plot (20−30 m total height) that was considered safe to climb. Ants were sampled in both day and night surveys with a combination of tuna baits and hand collection in ground and arboreal strata. Using the single-rope climbing technique, for each tree at every 3 m point from the ground to the highest part of the tree, we set five baited vials (∼1cm by 5cm), that were attached using tape and thumbtacks to the trunk, branches, and available epiphytes.

Ground baits were placed on the leaf litter or close to logs or other ground substrate. Baits were collected after three hours. During deployment and collection of baits, over a period of approximately one hour in each stratum, foraging ants were opportunistically hand collected using soft forceps and an aspirator. Day surveys began at 10:00 hr and night surveys began at 20:00 hr. Day and night surveys of the same trees were not conducted sequentially and were at least a day apart to allow resumption of normal ant activity in the tree. We note that our sampling of ground-foraging ants targets epigaeic species and is largely ineffectual for subterranean species. All voucher specimens were deposited in the ant collection held at the CSIROs Laboratory in Darwin, Australia.

### Measuring activity temperatures and ambient microclimate

Ant field activity temperatures were recorded as the surface temperature at the time of collection of every baited vial deployed every three metres from ground to canopy. Surface temperature was recorded using a handheld infrared temperature gun upon collection of every baited vial deployed every three metres from ground to canopy at each tree surveyed. Thus, for every ant capture in a baited vial we have a matched surface temperature for that specific surface location and time of day representing field active temperatures. To measure ambient microclimate temperature, we recorded temperature at 30-minute intervals using HOBO Pro v2 (U23-002) Temperature/Relative humidity data loggers placed at ground (∼ 0.5 m) and arboreal (∼ 20 m) habitats for two years from March 2019 – March 2021 at the 100 m a.s.l. and 1200 m a.s.l. sites. Data loggers were hung freely underneath a wide funnel (30 cm radius) to allow air flow while ensuring protection from extreme sun exposure which can degrade the equipment over long time periods and cause misleading spikes in temperature recordings.

### Thermal tolerance experiments

As soon as possible following collection, individual ants were tested for CT_min_ and CT_max_ in a digital dry bath using ramping assays as described by Leahy et al. (2022). Thermal tolerance data for daytime-collected ants has been reported previously (Leahy et al. 2022), whereas those for night-time collected ants are reported here for the first time.

### Data analysis

#### Quantifying thermal overlap

We aimed to quantify the extent to which temperatures overlapped between each combination of the vertical and temporal dimensions at the lowland and upland elevation site. Low thermal overlap would indicate that ant species are exposed to very different thermal conditions. For each elevation site and for each of the four-way combinations of ground, canopy, day, night (e.g., ground-day – ground-night, etc) we calculated a thermal overlap value as the overlap of two kernel density distributions using the function ‘overlap’ in the *overlapping* package (Pastore 2018). Note, the term canopy as a habitat is used in the microclimate analysis reflecting the placement of the thermal data logger in the canopy, whereas the term arboreal is used in downstream analyses to reflect that ants may be using the trunk of the tree below the canopy. Kernel density distributions consisted of recorded temperature values from the two years of continuous microclimate recordings. We plotted the density distributions using function ‘geom_density_ridges’ from the *ggridges* package (Wilke 2021).

#### Ant community composition and spatiotemporal activity

We compared community composition and species-specific spatiotemporal activity patterns amongst vertical habitats for each elevation site. For each plot at each elevation site, we pooled arboreal samples into one zone from 3 m to the highest vial deployment in each tree to compare to ground samples. Under this designation, the amount of time spent conducting hand collections was equal in ground and arboreal strata, but there was a difference in the number of baited vials deployed in ground and arboreal strata. We standardised species occurrence by total survey effort for ground and arboreal strata by dividing occurrences by the total number of vials deployed in each respective stratum.

We explored community composition patterns of ground versus arboreal and day versus night at each elevation site with multivariate ordination of standardised occurrence data (n = 57 species) constructed using Bray-Curtis dissimilarity. For lowland and upland sites, we tested the effect of vertical habitat (ground vs arboreal), time (day vs night), and their interaction on species composition using a PERMANOVA (using ‘adonis’ from *vegan* ver. 2.5; Oksanen *et al*. 2020) with “vertical habitat” and “day-night” as fixed factors and with “plot” as a random block factor and 999 permutations. Placing survey plot as a block factor accounted for our nested survey design (survey plot within elevation) and allowed the spatial variation in community structure created by turnover between individual plots to be accounted for in the model. We then standardised the data using the Wisconsin double standardization, which standardises the values in each row and column by the totals for that row and column respectively, thereby allowing easier computation and revealing more-subtle patterns of species composition (Legendre and Gallagher 2001). We tested for homogeneity of multivariate dispersions for our fixed factors using ‘betadisper’ in *vegan* ver 2.5 (Oksanen et al. 2020). We note that PERMANOVA with a balanced design, as in our models, is relatively robust to non-homogeneity of multivariate dispersions (Anderson and Walsh 2013). We used the percentage of explained variance (partial R^2^) to compare effect sizes between elevation sites.

To explore species-specific patterns of foraging activity in lowland and upland elevation sites, we selected the 21 common species with ≥ 4 total occurrences (100 m a.s.l., n = 13 species, 1200 m a.s.l., n = 8 species). This cut-off value was chosen to allow for the potential for a species to have been recorded in both strata and each time period (Houadria et al. 2015). We combined each species’ occurrences along vertical and temporal niche dimensions and then calculated a proportion of activity for each species in each of the four niche categories: arboreal day, ground day, arboreal night, ground night. Following Houadria *et al*. (2015), for each species we calculated the proportion of foraging activity in each elevation site by taking the mean plot-level relative frequency (taking the mean across the five plots sampled per site) for each of the categories (e.g., arboreal day) divided by the summed total frequency across the four categories for that species.

#### Field active thermal exposure and thermal tolerance

To explore the relationship between species activity, temperature exposure, and thermal tolerance we selected species for which we had upper and lower thermal tolerance data, ≥ 4 recorded sample occurrences, and a surface temperature recording from the baited vial collection (no species for this analysis occurred in both lowland and upland sites). We used the surface temperatures recorded for each species capture as their field foraging activity temperatures, we used temperature data collected from 2019 - 2020 (ground and canopy) as the species ambient microclimate temperature, and we used each species CT_min_ and CT_max_ as their thermal tolerance limits. These data are represented graphically, and implications discussed.

#### Thermal safety margins

We then asked: does having foraging flexibility to move into the cooler parts of the habitat improve thermal safety margins and does this vary between lowland and upland elevation sites. Thermal safety margins were calculated for seven species in the lowlands and five species in the uplands which were recorded foraging in more than one quadrant of the four-dimensional thermal space (quadrants could be = arboreal-day, arboreal-night, ground-day, ground-night). We calculated the thermal safety margin for the hottest quadrant of their total niche space (either daytime arboreal activity or daytime ground activity) as a baseline. We then calculated how much additional thermal safety margin would be afforded given they had the flexibility to move into a cooler part of their niche. Thermal safety margins were calculated as the species mean CT_max_ minus the mean foraging activity temperature (surface temperature), a proxy for operative temperature (T_e_) (Sinclair et al. 2016). Thermal safety margins could not be compared between the following foraging niches due to lack of species which demonstrated this combination of foraging activity, ground-day to ground-night at 100 m a.s.l., ground-day to arboreal-night at 1200 m a.s.l. These data are represented graphically, and implications discussed.

## Results

### Quantifying thermal overlap

In the lowlands, temperatures were more variable and therefore overlapped less between ground and arboreal habitats and between day and night compared to the uplands (Figure 1).

**Figure 1.**
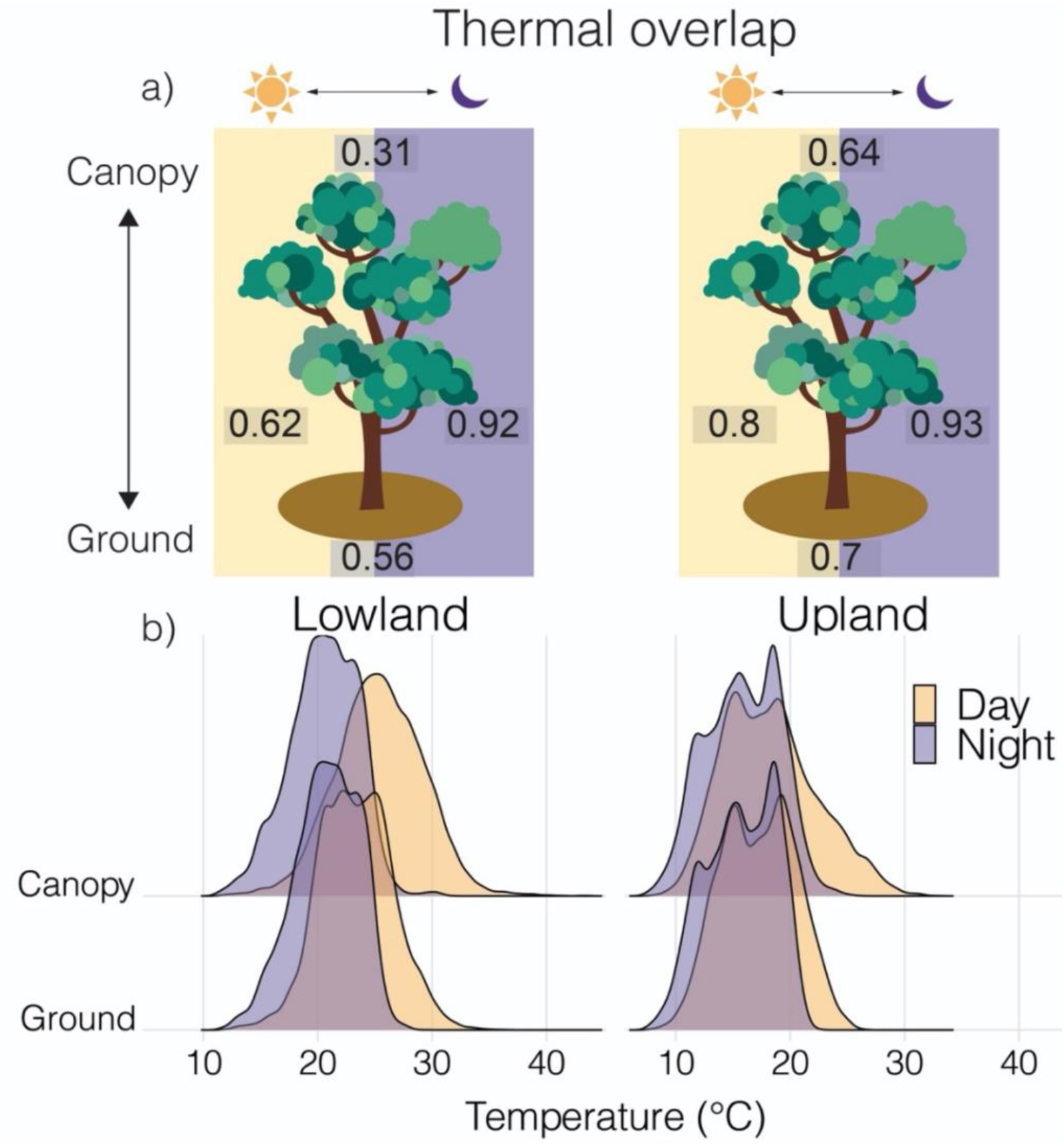
Thermal overlap along four-dimensional thermal niches - vertical (ground and canopy) and temporal (day and night) - and their different combinations at a lowland (100 m a.s.l.) and upland (1200 m a.s.l.) rainforest site in the Australian Wet Tropics Bioregion. Showing a) thermal overlap values (1 indicating complete thermal overlap) from b) kernel density distributions (y-axis) of microclimate air temperature recorded every half hour in each vertical position and elevation site for two years (March 2019-March 2021).

At both lowland and upland sites temperatures were more homogenised (there was a greater degree of thermal overlap) between ground and canopy at night (92% and 93% within lowland and upland, respectively) than in the day (62% and 80% within lowland and upland, respectively), and the thermal overlap between day and night-time temperatures was greater on the ground (56% and 70%, within lowland and upland respectively) compared to in the canopy (31% and 64%, within lowland and upland respectively).

### Spatiotemporal variation in ant species composition

There was markedly different species composition in ground and arboreal habitats at both elevations, but relatively little difference between day and night (Figure 2a-b, Table S1). There was no interactive effect of vertical habitat and time-period on species composition at either elevation (Table S1). Vertical habitat (independent of time period) explained 17% (pseudo-F_(1)_ = 3.96, p = 0.001) of variation in species composition at 100 m a.s.l. and 20% (pseudo-F_(1)_ = 4.29, p = 0.001) at 1200 m a.s.l. (Table S1). Time-period (independent of vertical habitat) explained 10% (pseudo-F_(1)_ = 2.27, p = 0.007) of variation in species composition at 100 m a.s.l. and 9% at 1200 m a.s.l. (pseudo-F_(1)_ =1.83, p = 0.05) (Table S1).

**Figure 2.**
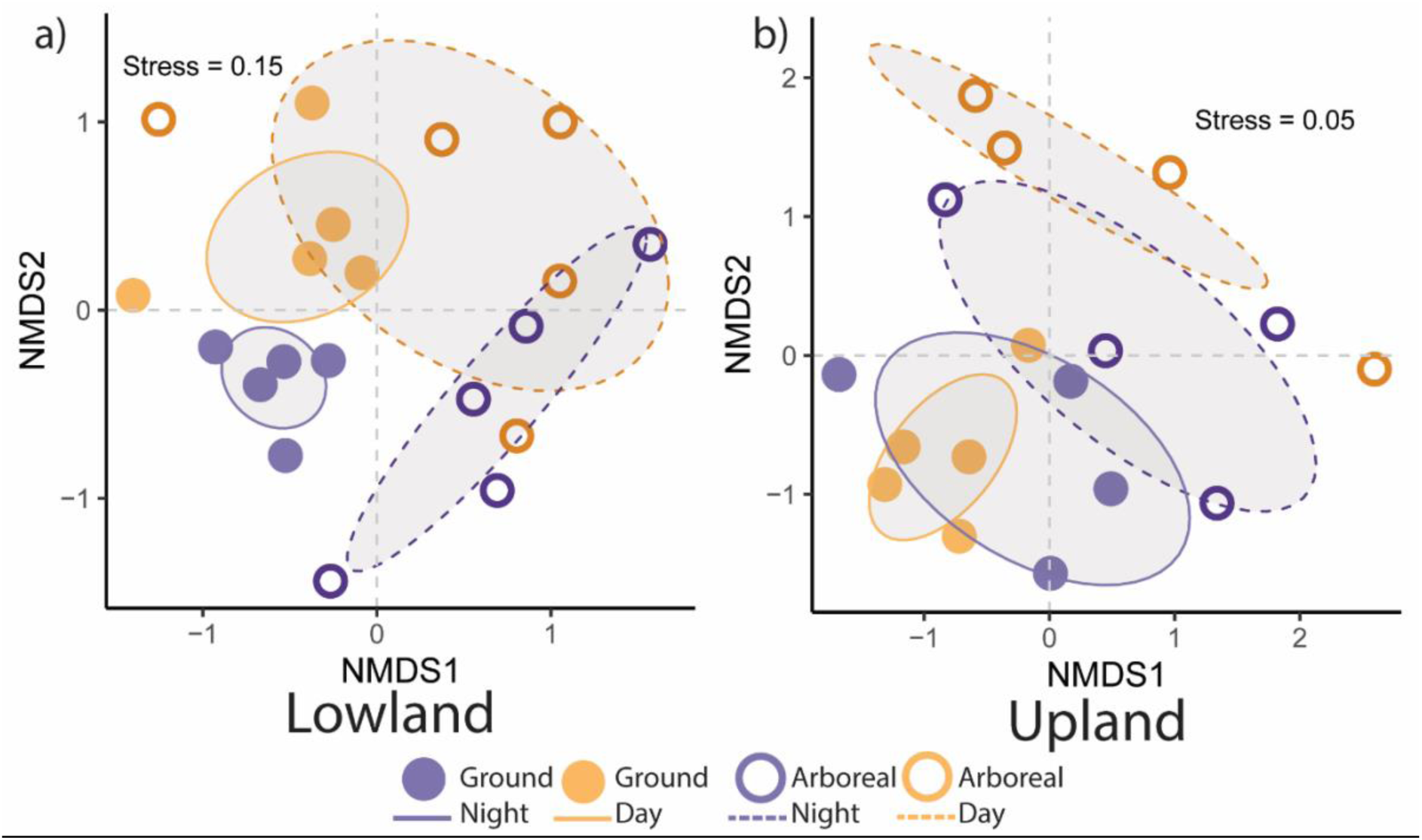
Spatiotemporal ant community structure showing NMDS ordination of ant species composition in ground and arboreal habitats in day and night-time surveys at two elevations in the Australian Wet Tropics Bioregion. A) 100 m a.s.l. (n = 42 species), B) 1200 m a.s.l. (n = 18 species). Polygons show grouping of surveys/sites where further separation between polygons represents greater differences in species composition.

### Vertical and temporal foraging activity

At the temporal scale, species analysed for foraging activity patterns (those with ≥ 4 total occurrences) were nearly all captured in both day and night samples: 77% and 87.5% of species at the 100 and 1200 m a.s.l. sites, respectively (Figure 3). Most of these species were also recorded in both ground and arboreal habitats at the 100 m a.s.l. (60%) site (Figure 3a). At 1200 m a.s.l., however, about half the species were vertically specialised, recorded either solely on the ground or in arboreal habitat (Figure 3b).

**Figure 3:**
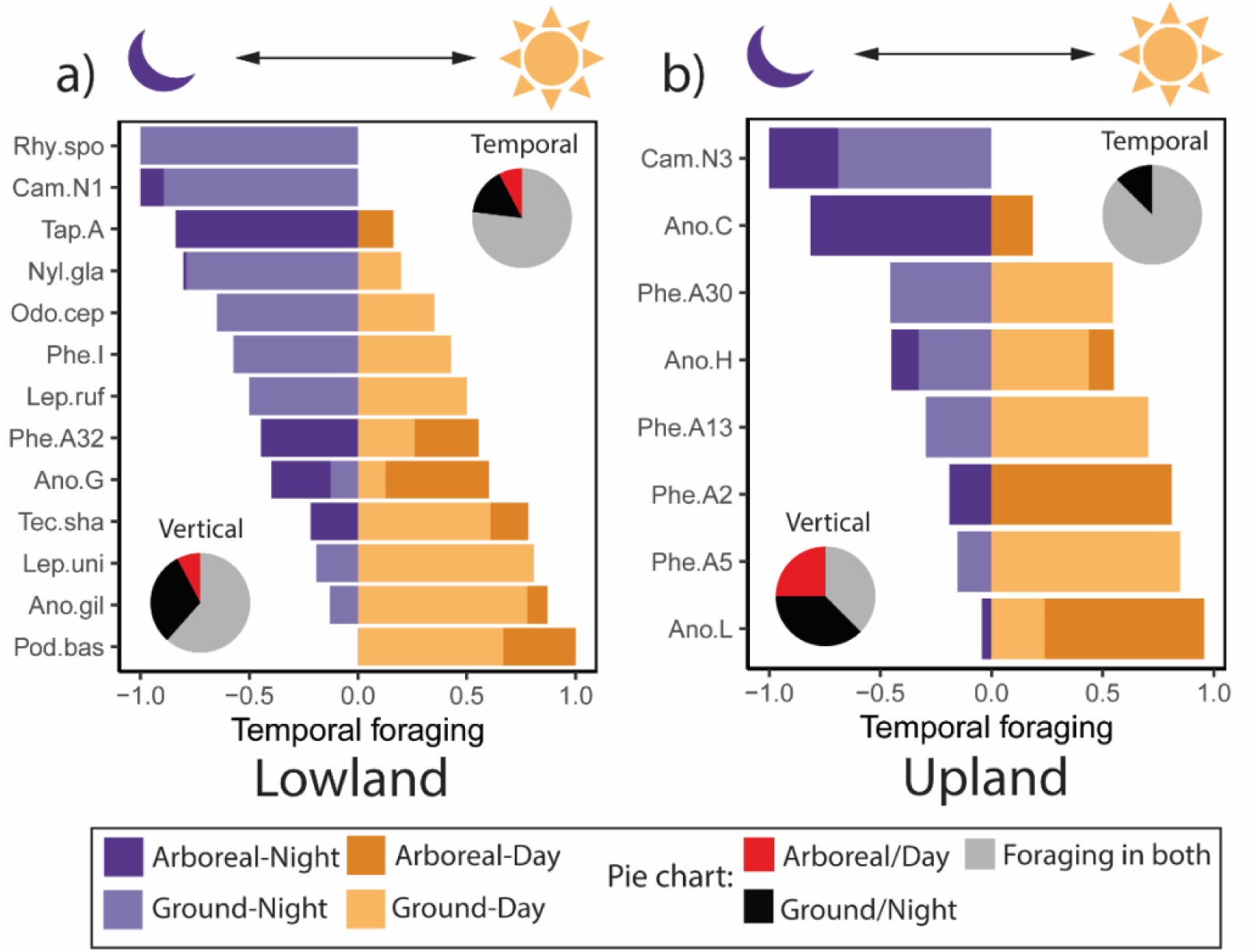
Temporal and vertical activity of ants based on species relative frequencies in surveys conducted in ground and arboreal habitats in the day or night along an elevation gradient in the Australian Wet Tropics Bioregion. Showing common species with four or more occurrences at a) 100 m a.s.l. b) 1200 m a.s.l. Bar charts show activity frequency, where negative values indicate night-time activity, and positive values indicate daytime activity.

Within each bar, dark colours indicate arboreal activity, and light colours indicate ground activity. Pie charts show temporal and vertical separately and are representing the proportion of species with activity recorded in both niche categories and activity only recorded in one niche category. Abbreviated names are first three letters of genus and species. Full species names are shown in Table S2.

### Activity, thermal exposure, and thermal tolerance

Of the 16 species for which we had thermal limit assays, we found that most were foraging at temperatures well below both their CT_max_ values and maximum ambient air temperatures for their elevation site. For lowland ant community, CT_min_ ranged from 6.6–9.8 °C, and CT_max_ from 42–49.3 °C (Figure 4a). Maximum air temperature reached 43.6 °C in the mid to high canopy surpassing the CT_max_ of three of the eight species from that community (Figure 4a, Table S2), however, daytime field active temperatures were generally below 30 °C even in the high canopy. For the upland ant community, CT_min_ ranged from 3.6–5.7 °C and CT_max_ from 36.9–47.4 °C (Figure 4b). Maximum air temperatures were below the lowest CT_max_ by ∼10 °C on the ground and ∼4 °C in the canopy in the upland site (Figure 4b). In both sites, field active daytime foraging temperatures were ∼8-fold more variable across the vertical gradient compared to nighttime foraging temperatures (daytime field active temperatures: σ^2^ = 15.5 °C (lowland) and σ^2^ = 17.4 °C (upland), nighttime field active temperatures: σ^2^ = 2.4 °C (lowland), σ^2^ = 2.2 °C (upland)).

**Figure 4:**
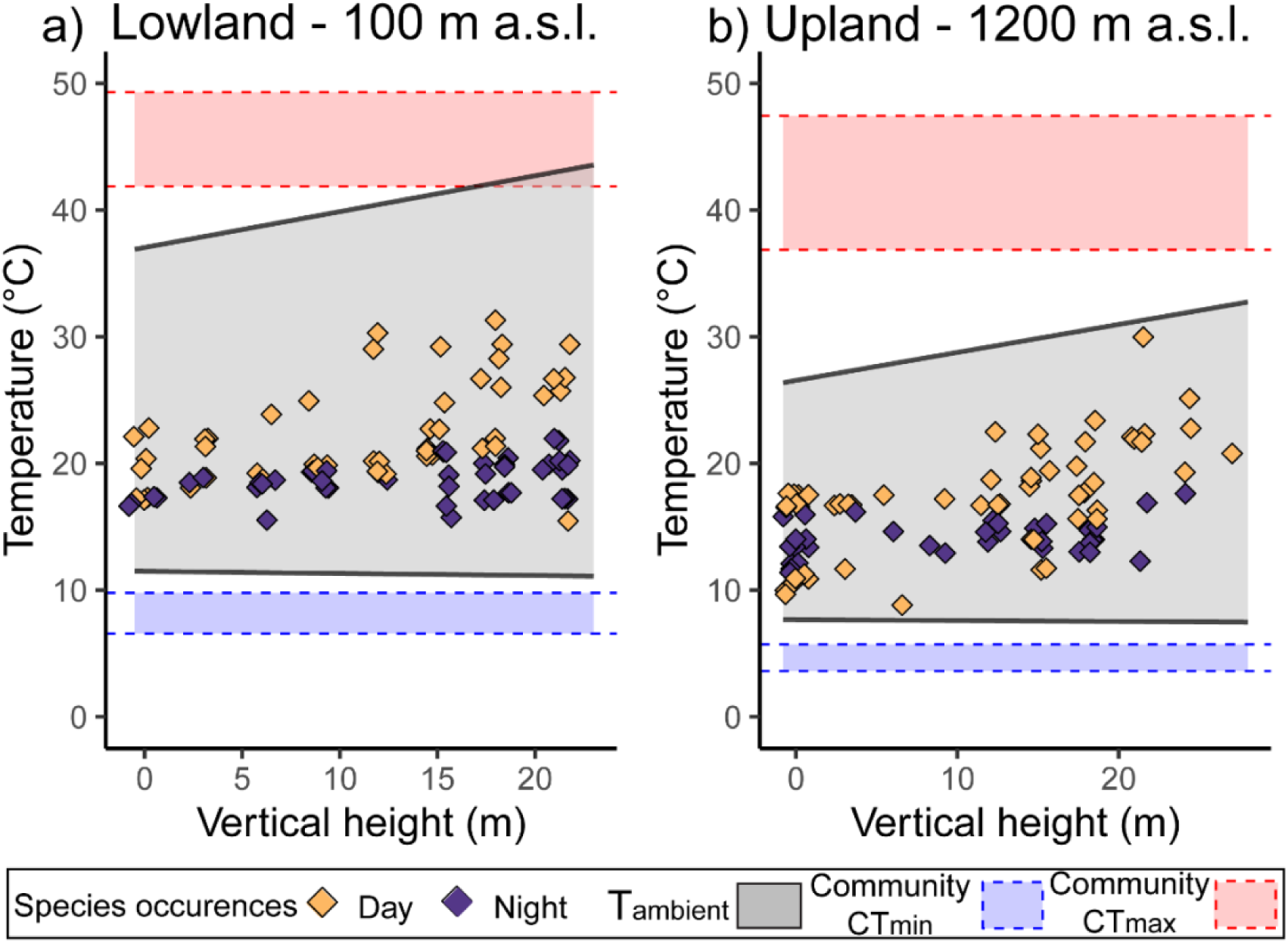
Field active surface temperatures in day and night for each ant species occurrence (diamonds) in a) lowland (100 m a.s.l.) and b) upland (1200 m a.s.l.) rainforest in relation to two years of ambient temperature (T_a_) microclimate recordings (grey shaded area, black lines) and community thermal tolerance limits (CT_min_ = blue, CT_max_ = red, shaded area, dashed lines). Showing common species with four or more survey occurrences and thermal tolerance data (lowland: n = 8 species, upland: n = 8 species). Microclimate temperatures show average minimum and maximum ambient temperature recorded in ground and canopy habitats over two years (2019 – 2021). Ant surveys conducted along vertical (ground to canopy) and in day (10:00 – 16:00 hrs) and night (19:00 – 22:30 hrs) time periods over ten days.

### Spatiotemporal flexibility and thermal safety margins

Foraging in cooler parts of the vertical and temporal thermal landscape increased thermal safety margins of ant communities at both elevations, but increases were more dramatic in the lowlands. Using either ground-day or arboreal-day as the hottest niche baselines, thermal safety margin increased by an average of 0.84 °C (± 0.73 SE) for ground-day to arboreal-night shifts, 4.39 °C (± 1.4 SE) for arboreal-day to ground-day shifts, and 6.7 °C (± 1.35 SE) for arboreal-day to arboreal-night shifts in the lowlands (Figure 5a). In the uplands, increases in thermal safety margins were similar for each niche shift from baselines: averaging 1.8 °C (± 1 SE) increase for ground-day to ground-night, and an average of 2.1 °C increase for arboreal-day to ground-day (± 0.28 SE) and arboreal-day to arboreal-night (± 2.1 SE) (Figure 5b).

**Figure 5:**
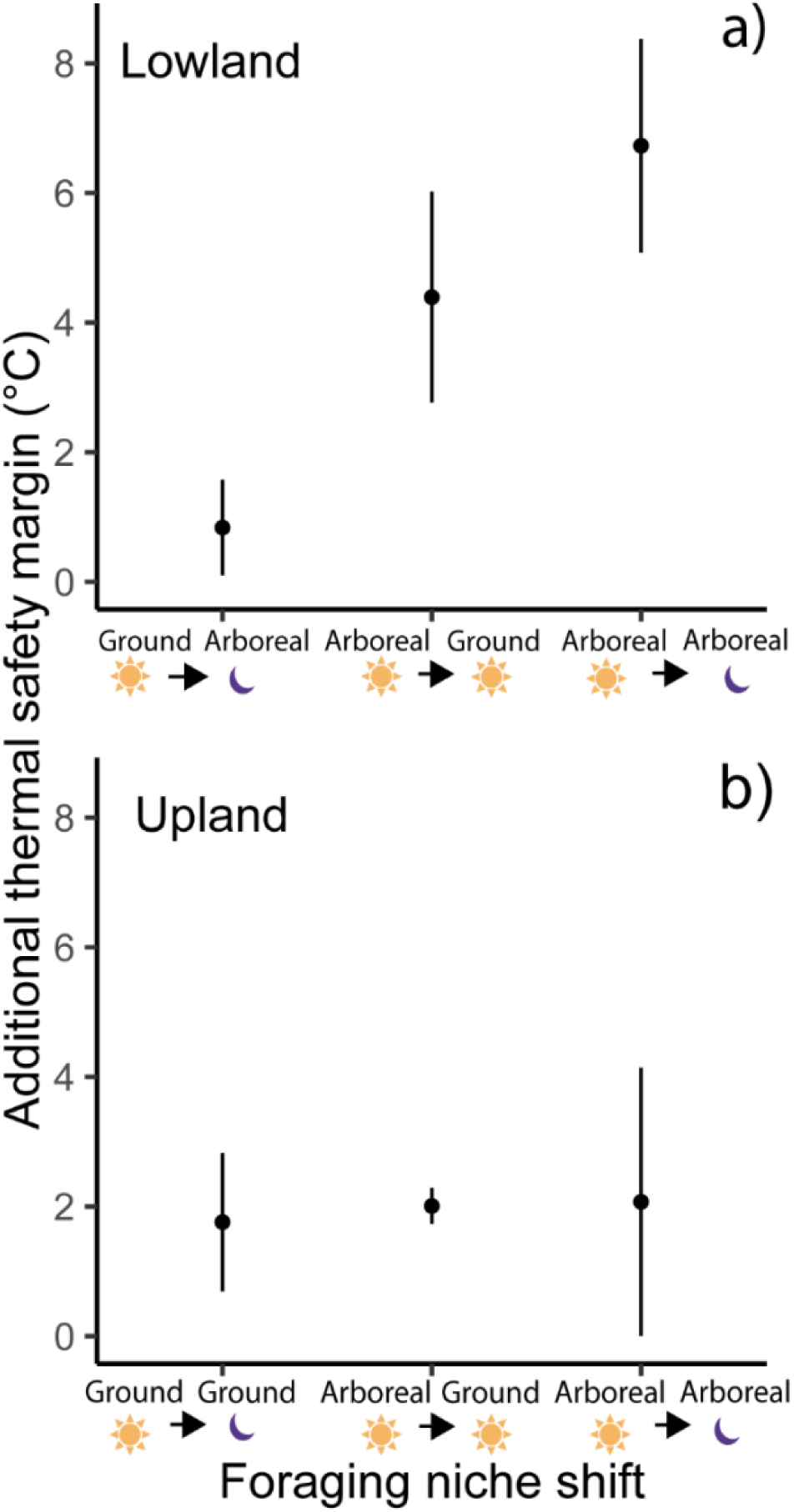
Rainforest ants that inhabit warmer spatiotemporal niches increase their thermal safety margins by using flexible foraging behaviour to move into cooler habitats and activity times at a) lowland (100 m a.s.l.) and b) upland (1200 m a.s.l.) sites. Foraging niche shifts on the x-axis shows moving from the spatiotemporal niche on the left (representing the hotter parts of the niche as a baseline), to the niche on the right and the associated mean increase (± SE) in thermal safety margin. Sun symbols represents diurnal activity, moon symbols represents nocturnal activity.

## Discussion

Despite tropical rainforests typically being characterised as climatically stable, the vertical gradient and diel cycle produce a four-dimensional thermal landscape at the local habitat scale. At the Carbine range, we found the thermal landscape to span more than 30°C range of temperatures in the lowlands and a 25°C range in the uplands. We describe species activity and thermal ecology along these spatiotemporal thermal gradients at a warm low and cool high elevation and explore the capacity for rainforest ants to buffer climate change *in situ* through existing flexibility in the location and timing of activity. We found that a large proportion of species at lowland and upland elevations have the capacity to be active in the cooler hours of the night and towards the (cooler) ground. This flexibility substantially increased thermal safety margins, particular in the lowlands, thereby reducing thermal exposure risk.

Overall, the upland thermal landscape was more homogenous compared to the lowlands (see Figure 1b) with higher overlap of thermal profiles across vertical and temporal gradients. Similarly, in a neotropical forest, temperature profiles of open habitat (comparable to the hot and variable canopy) and forested habitat became more similar with increasing elevation (Kumar and O’Donnell 2009). At lowland and upland elevations, ground and canopy thermal profiles were divergent during the day but became homogenised at night (Figure 1). In addition, ant field active surface temperatures were ∼8-fold more variable for diurnal foragers than for nocturnal foragers (Figure 4), emphasising the distinct thermal profile of canopy environments.

We found stronger vertical stratification relative to temporal differentiation in ant communities at both elevation sites, but this was more pronounced in the upland site, where vertical habitat explained twice as much variation in community composition than did temporal niche. Our results echo Kaspari and Weiser’s (2000) study that found little distinction between diurnal and nocturnal ant communities but strong vertical stratification. Similarly, in another study from the Australian Wet Tropics, Bluthgen et al. (2004) reported that most lowland canopy ants were foraging both day and night. In our study, although vertical stratification was stronger than temporal differences in activity, we did not find very high rates of vertical specialisation per se compared to other tropical realms. For example, in the Neotropics, canopy and litter ant communities are highly vertically specialised with no shared species across the domains (Yanoviak and Kaspari 2000). Here, vertical habitat alone explained no more than 20% of community composition patterns, and the majority of ants at every elevation foraged across all four dimensions of the spatiotemporal landscape (see pie-charts, Figure 3).

Flexible temporal activity in ants – where species are active both day and night or switch seasonally between day and night – has been reported in a wide range of ecosystems such as; temperate forests and grasslands (Fellers 1989, Albrecht and Gotelli 2001, Stuble et al. 2013, Żmihorski and Ślipinski 2016), semi-arid zones of south-America and Australia (Briese and Macauley 1980, Andersen 1983, Bestelmeyer 2000), desert (Whitford 1978, Heatwole and Muir 1991), neotropical and paleotropical rainforests (Yamane et al. 1996, Kaspari and Weiser 2000, Blüthgen et al. 2004, Hashimoto et al. 2010, Tanaka et al. 2010, Houadria et al. 2015, Yusah et al. 2018, Grevé et al. 2019, Houadria and Menzel 2020), Mediterranean grasslands (Cros et al. 1997, Cerdá et al. 1998a, Cerdá et al. 1998b, Retana and Cerdá 2000), and the Caatinga xeric shrublands of Brazil (Silva et al. 2019). This wide geographic spread shows that flexible temporal activity is a common feature of ant ecology, but the mechanistic basis of this strategy across thermal gradients has not been investigated in detail.

Against our prediction, temporal flexibility in activity was surprisingly strong in the uplands, given that ants generally prefer foraging under warmer conditions (Dunn et al. 2009). This behaviour may confer benefits by increasing the time window of food acquisition, since upland systems are less productive with scarcer resources and ants from wet and cool sites tend to have lower metabolic rates (Lasmar et al. 2021, Leahy et al. 2025a). Bishop et al. (2015) found that specialised trait combinations were lost from ant assemblages at high elevation such that ants tended to become dietary generalists and have no preference for open or closed habitats. Relatively low interspecific competition as a result of low species richness might also be a factor contributing to lower temporal niche specialisation in uplands (Camarota et al. 2016).

The widespread flexibility in activity that we found for both vertical and temporal niches at both elevation sites indicates an ecological strategy with important implications for resilience to climate change (Woods et al. 2015, Bonebrake et al. 2020, Enriquez-Urzelai et al. 2020, Gibb et al. 2023). The field active (surface) temperatures we recorded during our surveys were often well below species upper thermal limits, which provides a buffer between current operative temperatures and maximum lethal temperatures (Huey et al. 2012, Sinclair et al. 2016). Maximum ambient temperatures in the canopy did at times surpass the upper thermal limits of some species in the lowlands. However, as predicted, species with flexibility in activity times and/or vertical activity could substantially decrease this risk by moving into cooler parts of their thermal landscape (Figure 5).

Lowland canopy active species increased their thermal safety margins by an average of 4.4 °C by foraging on the ground and 6.7 °C by selecting to forage at nighttime. Under future climate change, this pre-existing behavioural flexibility could make a substantial difference during heatwave events (Moritz and Agudo 2013, Sunday et al. 2014, de la Fuente et al. 2025). This could allow ant communities to mitigate climate risk *in situ*, as demonstrated for mammal species in a recent global analysis: obligate diurnal and nocturnal specialists were more than twice as likely to have already shown a geographic range shift response to climate change than mammals with flexible activity times (Román-Palacios and Wiens 2020). Conversely, lowland canopy specialists with an obligate diurnal activity have the highest levels of thermal exposure and the least capacity for reducing exposure to extreme temperatures. Interestingly, we found very few ant species that fit that description in our Australian Wet Tropics sites. One hypothesis is that diel specialisation towards diurnal foraging has been selected against in rainforest ants, this warrants a more geographically widespread investigation across tropical bioregions (Williams et al. 2009, Wong and Didham 2024).

Several caveats of our findings must be noted. Firstly, our thermal tests took place in the cooler dry season and temperate forest ants have been known to shift their upper thermal limit with season (Bujan et al. 2020). We do not know if tropical forest ants undergo seasonal plasticity in upper thermal limits, and this should be an area for further investigation. Secondly, we did not survey for ants at every hour of the day, and we do not have longer-term data on field active temperatures that ants might experience over a full year. Therefore, ants might be active at hotter temperatures than we have recorded here. Finally, the upland community was restricted in phylogenetic diversity consisting mostly of species from the *Pheidole* and *Anonychomyrma* genera. These genera were also present in the lowlands, providing a good comparison of microclimatic exposure and sensitivity of these dominant genera, however, as the lowland and upland sites were not replicated across multiple mountain ranges, we urge further exploration and confirmation of the patterns reported here in other tropical systems.

Our findings have broad implications for rainforest biodiversity under climate change. In general, small-bodied ectotherms with any degree of vertical and/or diel niche breadth could be utilising the four-dimensional thermal landscape of the rainforest to mitigate temperature increases, however, the extent of spatiotemporal specialisation is known for very few tropical taxa (Gaston 2019). Behavioural buffering of temperatures may come with associated costs to foraging success and fitness, and could forestall physiological adaptation to increased temperatures (Buckley et al. 2015, Muñoz and Losos 2018). Quantifying these costs, particularly incorporating ecological traits, energetics, and life history syndromes (Leahy et al. 2025b, Riskas et al. 2026), is the next research step to further understand this mechanism of change climate mitigation. We conclude that species with specialised microhabitat and activity niches that are restricted to the hottest parts of the rainforest are most vulnerable to climate change (Colwell et al. 2008, Corlett 2011, Diamond et al. 2012, Senior et al. 2019). This should raise concern given the extraordinary invertebrate diversity in the canopies of lowland tropical rainforest (Novotný and Basset 2000, Basset et al. 2003, Ozanne et al. 2003, Basset et al. 2012, Ashton et al. 2016).

## Acknowledgements and statement of inclusion

We thank the Jabalbina Yalanji Aboriginal Corporation and acknowledge the traditional owners of the Eastern Kuku Yalanji and Western Yalanji on whose lands this work was conducted. We thank our funders: The Explorer’s Club, Wet Tropics Management Authority, Skyrail Rainforest Foundation, Holsworth Wildlife Research Endowment – Equity Trustees Charitable Foundation. LL was supported by a PhD scholarship from the Australian Government. Field research conducted under Permit WITK15811415 issued by the Queensland Government of Australia. Our study brings together authors from a number of different countries, including scientists based in the country where the study was carried out. Whenever relevant, literature published by scientists from the region was cited. Field work was conducted with permission and a welcome to country from the traditional custodians of the Mossman Gorge site. A report on the outcomes of the research and ant collection data with photos were provided to the Jabalbina Yalanji Aboriginal Corporation upon completion of the work.

## Statement of contribution of authors

The study was conceived by LL, BRS, SEW and ANA; LL collected all data; ANA identified ant species and managed the ant collection; LL conducted all analyses and wrote the first draft of the manuscript; all authors contributed critically to subsequent drafts and gave final approval for publication.

