## Supporting Information 1 for "The nightshift lowdown: can ants buffer climate change through shifts in vertical and temporal activity?"

**Table S1.** The effect of vertical habitat and time-period on community composition of ants along an elevation gradient in the Australian Wet Tropics Bioregion. Results of PERMANOVA based on species relative frequency in ground and arboreal surveys conducted in day and night. With “Vertical habitat” and “Time-period” and their interaction as fixed factors and “plot” as a random block factor and using Bray-Curtis dissimilarity. Significance threshold: p < 0.05.

| **Elevation (m a.s.l.)** | **Vertical** (df = 1) |  |  | **Time-period** (df =1) |  |  | **Vertical:Time-period** (df = 1) |  |  | **Residuals** (df) |
| --- | --- | --- | --- | --- | --- | --- | --- | --- | --- | --- |
|  | Pseudo F | *p* | % Explained variance | Pseudo F | *p* | % Explained variance | Pseudo F | *p* | % Explained variance | % Explained variance |
| 100 | 3.96 | 0.001 | 17 | 2.27 | 0.007 | 10 | 1.22 | 0.187 | 5 | 0.68 (16) |
| 1200 | 4.29 | 0.001 | 20 | 1.83 | 0.05 | 9 | 1.0 | 0.443 | 5 | 0.66 (14) |

**Table S2.** Site elevation, species, number of plot and sample occurrences, relative frequency of occurrence in arboreal and ground during daytime and night-time, and thermal tolerance (n = 1–3 colonies/species). Showing species with four or more sample occurrences.

| Site | Species |  | | Activity | | | | | Thermal traits | |
| --- | --- | --- | --- | --- | --- | --- | --- | --- | --- | --- |
|  |  | Plot (n) | Sample (n) | Arboreal Day | Arboreal Night | Ground Day | Ground Night | Mean CT_min_ ± SD (ºC) | | Mean CT_max_ ± SD (ºC) |
| 100 | *Anonychomyrma gilberti* | 4 | 12 | 0.09 | 0.00 | 0.78 | 0.13 | 6.6 ± 1.36 | | 43.43 ± 2.78 |
|  | *Anonychomyrma* sp. G | 2 | 38 | 0.48 | 0.27 | 0.13 | 0.13 | 6.58 ± 1.17 | | 45.97 ± 1.51 |
|  | *Camponotus* sp. N1 | 5 | 11 | 0.00 | 0.11 | 0.00 | 0.89 | 6.8 ± 0.41 | | 41.87 ± 2.88 |
|  | *Leptomyrmex ruficeps* | 2 | 4 | 0.00 | 0.00 | 0.50 | 0.50 | 6.15 ± 2.27 | | 38 ± 5.13 |
|  | *Leptomyrmex unicolor* | 4 | 5 | 0.00 | 0.00 | 0.81 | 0.19 | 10.4 ± 0.76 | | 41.6 |
|  | *Nylanderia glabrior* | 5 | 11 | 0.00 | 0.01 | 0.20 | 0.79 | 7.23 ± 0.7 | | 41.93 ± 2.4 |
|  | *Odontomachus cephalotes* | 5 | 9 | 0.00 | 0.00 | 0.35 | 0.65 | 11.86 ± 1.23 | | 41.45 ± 1.45 |
|  | *Pheidole* sp. A32 | 3 | 21 | 0.29 | 0.45 | 0.26 | 0.00 | 8.27 ± 1.02 | | 42.03 ± 1.52 |
|  | *Pheidole* sp. I | 3 | 5 | 0.00 | 0.00 | 0.43 | 0.57 | 8.74 ±1.1 | | 40.49 ± 2.52 |
|  | *Podomyrma basalis* | 2 | 4 | 0.33 | 0.00 | 0.67 | 0.00 | 9.78 ± 1.8 | | 49.3 ± 3.25 |
|  | *Rhytidoponera spoliata* | 4 | 4 | 0.00 | 0.00 | 0.00 | 1.00 | 6.41 ± 0.88 | | 42.49 ± 0.83 |
|  | *Tapinoma* sp. A | 2 | 6 | 0.16 | 0.84 | 0.00 | 0.00 | 7.32 ± 2.01 | | 45.1 ± 0.14 |
|  | *Technomyrmex shattucki* | 2 | 11 | 0.17 | 0.22 | 0.61 | 0.00 | 7.19 ± 1.04 | | 44.48 ± 1.83 |
| 1200 | *Anonychomyrma* sp. C | 1 | 9 | 0.19 | 0.81 | 0.00 | 0.00 | - | | - |
|  | *Anonychomyrma* sp. H | 3 | 26 | 0.11 | 0.12 | 0.44 | 0.33 | 3.84 ± 1.14 | | 47.44 ± 2.62 |
|  | *Anonychomyrma* sp. L | 1 | 23 | 0.72 | 0.04 | 0.24 | 0.00 | 3.61 ± 0.2 | | 42.48 ± 1.12 |
|  | *Camponotus* sp. N3 | 2 | 8 | 0.00 | 0.31 | 0.00 | 0.69 | 4.01 ± 0.78 | | 40.76 ± 0.52 |
|  | *Pheidole* sp. A13 | 4 | 8 | 0.00 | 0.00 | 0.71 | 0.29 | 5.72 ± 0.93 | | 39.48 ± 1.14 |
|  | *Pheidole* sp. A2 | 3 | 11 | 0.81 | 0.19 | 0.00 | 0.00 | 5.47 ± 0.79 | | 38.9 ± 0.93 |
|  | *Pheidole* sp. A30 | 5 | 10 | 0.00 | 0.00 | 0.55 | 0.45 | 4.2 ± 0.19 | | 36.86 ± 0.4 |
|  | *Pheidole* sp. A5 | 4 | 8 | 0.00 | 0.00 | 0.85 | 0.15 | 5.18 ± 0.29 | | 37.87 ± 1.1 |
